# Epigenetic clock and methylation studies in cats

**DOI:** 10.1101/2020.09.06.284877

**Authors:** Ken Raj, Balazs Szladovits, Amin Haghani, Joseph A. Zoller, Caesar Z. Li, Steve Horvath

## Abstract

Human DNA methylation profiles have been used successfully to develop highly accurate biomarkers of aging (“epigenetic clocks”). Although these human epigenetic clocks are not immediately applicable to all species of the animal kingdom, the principles underpinning them appear to be conserved even in animals that are evolutionarily far removed from humans. This is exemplified by recent development of epigenetic clocks for mice and other mammalian species. Here, we describe epigenetic clocks for the domestic cat (*Felis catus*), based on methylation profiles of CpGs with flanking DNA sequences that are highly conserved between multiple mammalian species. Methylation levels of these CpGs are measured using a custom-designed Infinium array (HorvathMammalMethylChip40). From these, we present 3 epigenetic clocks for cats; of which, one applies only to blood samples from cats, while the remaining two dual-species human-cat clocks apply both to cats and humans. As the rate of human epigenetic ageing is associated with a host of health conditions and pathologies, it is expected that these epigenetic clocks for cats would do likewise and possess the potential to be further developed for monitoring feline health as well as being used for identifying and validating anti-aging interventions.

## Introduction

Most owners of domestic cats lament the short lifespan of these widely popular pets. The oldest recorded age of a cat is 30 years, however most cats succumb to diseases and die before they are 20 years old. Age is undoubtedly the biggest risk factor for a vast majority of diseases in animals, and cats are no exception. Interventions to retard or stop aging are being sought, and some are already being tested in humans and rodents. These investigations however, have yet to be extended to cats even though they share similar environments and living conditions with their human owners. Identification of environmental factors and living conditions that affect ageing, as well as potential mitigation measures, can be achieved by proxy, with cats. A prerequisite for such investigations however, is a highly accurate set of biomarkers of aging for both, cats and humans. While there is a rich literature on human epigenetic clocks for various individual and multiple tissues ^1-4^, we are not aware of any existing epigenetic clocks for cats. The human epigenetic clocks have already found many biomedical applications including the measure of biological age in human anti-ageing clinical trials ^1,5^. This has instigated the development of similar clocks for other mammals such as mice and dogs ^6-12^. Here, we aimed to develop and evaluate epigenetic clocks for cats, as such biomarkers are necessary for translating promising anti-aging interventions from humans to cats and *vice versa*, and to also provide the possibility of using the epigenetic ageing rate of cats to inform on feline health; for which a quantitative measure is presently unavailable. Specifically, we present here, DNA methylation-based biomarkers (epigenetic clocks) of age for blood from cats.

It was known for a long while that the degree of cellular DNA methylation alters with age ^13-15^. The significance and specificity of these alterations remained a source of speculation until the development of an array-based technology that permitted the simultaneous quantification of methylation levels of specific CpG positions on the human genome. With this technical advancement, came the opportunity and insight to combine age-related methylation changes of multiple DNA loci to develop a highly accurate age-estimator for all human tissues ^1,2,4^. For example, the human pan-tissue clock combines the weighted average of methylation levels of 353 CpGs into an age estimate that is referred to as DNAm age or epigenetic age ^16^. As would be expected of such an age-estimator, its prediction of epigenetic age corresponds closely to chronological age. What is significantly more important however, is the finding that the discrepancy between epigenetic age and chronological age (which is termed “epigenetic age acceleration”) is predictive of multiple health conditions, even after adjusting for a variety of known risk factors ^17-22^. Specifically, epigenetic age acceleration is associated with, but not limited to cognitive and physical functioning ^23^, Alzheimer’s disease ^24^, centenarian status ^21,25^, Down syndrome ^26^, progeria ^27,28^, HIV infection ^29^, Huntington’s disease ^30^, obesity ^31^, menopause ^32^. Epigenetic age is also predictive of mortality even after adjusting for known risk factors such as age, sex, smoking status, and other risk factors ^17-22^. Collectively, the evidence is compelling that epigenetic age is an indicator of biological age ^33-35^.

We previously demonstrated that the human pan-tissue clock, which was trained with only human DNA methylation profiles can be applied directly to chimpanzees, ^16^ but it loses utility for other animals as a results of evolutionary genome sequence divergence. Recently, others have constructed epigenetic clocks for mice and successfully validated them with benchmark longevity interventions including calorie restriction and growth hormone receptor knockouts ^6-11^. Overall, these independent initiatives indicate that the underlying biological principle of epigenetic clocks is shared between members of different species within the mammalian class, and that it is possible and feasible to extend the development of epigenetic clocks to other mammalian species. Our current study pursued the following aims. First, to develop a DNA-methylation-based estimator of chronological age across the entire lifespan of cats and humans (dual species clocks). Second, to characterize age-related changes in methylation levels in cats. Third, to test the performance of these cat epigenetic clocks on four different species of the mammalian class: guinea pigs, rabbits, ferrets, and alpacas.

## Results

We generated DNA methylation profiles from n=130 blood samples from cats of different breeds, whose ages ranged from 0.21 to 20.9 years **(Table 1** and **Table 2)**. Cat breeds did not correspond to distinct clusters in the unsupervised hierarchical clustering analysis (color band in **Figure 1**). Similarly, a supervised machine learning analysis (based on the random forest predictor) did not lend itself for predicting cat breed.

**Table 1.** Description of blood methylation data from 5 different species. N=Total number of samples per species. Number of females. Age: mean, minimum and maximum.

| Species | Latin Name | N | No. Female | Mean Age | Min. Age | Max. Age |
| --- | --- | --- | --- | --- | --- | --- |
| Domestic cat | *Felis catus* | 128 | 58 | 6.93 | 0.211 | 20.9 |
| Guinea pig | *Cavia porcellus* | 2 | 1 | 1.67 | 0.917 | 2.42 |
| Ferret | *Mustela putorius furo* | 2 | 2 | 2.67 | 0.5 | 4.83 |
| European rabbit | *Oryctolagus cuniculus* | 5 | 0 | 3.02 | 1 | 4.5 |
| Alpaca | *Vicugna pacos* | 5 | 3 | 7.93 | 1.67 | 15 |

**Table 2.** Description of cat breeds used for the development of cat clocks.

| Cat Breed | Number |
| --- | --- |
| Abyssinian | 1 |
| Bengal | 4 |
| Birman | 3 |
| British Blue | 6 |
| British Shorthair | 8 |
| Burmese | 6 |
| Cornish Rex | 1 |
| Devon Rex | 1 |
| Domestic Long Hair | 11 |
| Domestic Medium Hair | 2 |
| Domestic Short Hair | 57 |
| European Short Hair | 1 |
| Exotic | 2 |
| Korat | 1 |
| Main Coon | 6 |
| Norwegian Forest Cat | 1 |
| Persian | 3 |
| Ragdoll | 6 |
| Russian Blue | 1 |
| Scottish Fold | 1 |
| Siamese | 3 |
| Siberian Forest Cat | 3 |
| Sphynx | 2 |

**Figure 1.**
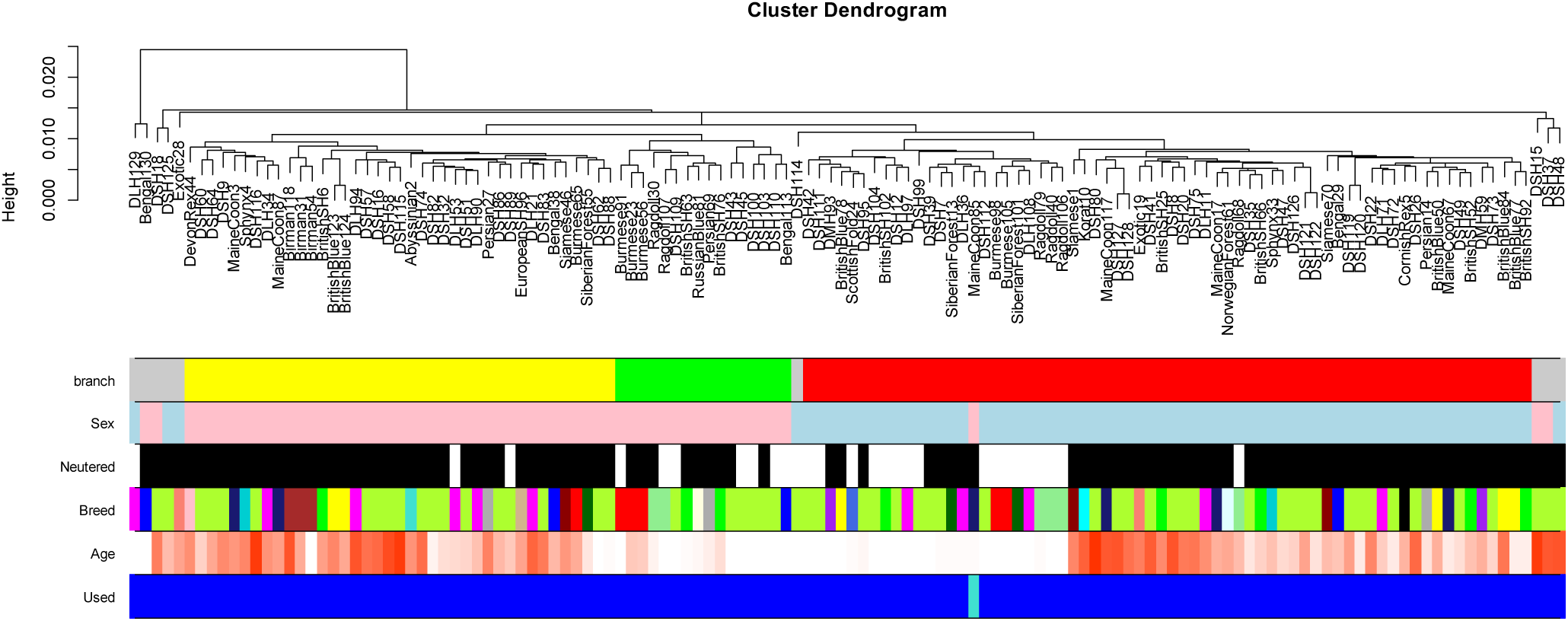
Unsupervised hierarchical clustering of blood samples from cats. Average linkage hierarchical clustering based on the inter-array correlation coefficient (Pearson correlation). The low height values (y-axis) indicate high inter-array correlations (R>0.97) and high quality. The cluster branches (first color band) correspond to sex (second color band, pink=female). Third color band: neutered (black) versus intact (white). Fourth color band encodes cat breeds: magenta=domestic long hair, greenyellow=domestic shorthair, blue=bengal, lightgreen=ragdoll, green=British shorthair, darkred=Siamese. One sample (last color band) was omitted from the analysis because its sex did not match that of its neighboring samples.

Most of the blood samples were from neutered cats: 50 from neutered females, 51 from neutered males, 9 from intact females and 19 from intact males. Their DNA methylation profiles clustered by sex (**Figure 1**). One of the cat samples was removed from the analysis because its sex did not match the clustering pattern. A subsequent random forest analysis of sex led to perfect (out-of-bag, OOB) estimates of accuracy (zero misclassifications). Neutered status was strongly confounded with age: 25 out of 28 intact animals were younger than 0.8 years. A random forest prediction analysis of neutered status led to a high OOB error rate (>60% error rate) when restricting the analysis to the young samples.

We also generated blood DNA methylation profiles from 4 additional mammalian species: two guinea pigs, five rabbits, two ferrets, and five alpacas (**Table 1**). As would be expected, the methylation profiles of these were clearly clustered by species (**Supplementary Figure 1**).

## Epigenetic clocks

We developed three epigenetic clocks for cats that vary with regards to two features: species and measure of age. The basic cat clock, which is constituted by 34 CpGs, was trained on cat blood DNA methylation profiles, while the dual-species (human-cat) epigenetic clocks were trained using cat and human DNA methylation data. The resulting two human-cat clocks mutually differ by way of age measurement. One estimates *chronological age*s of cats and humans (in units of years) based on methylation profiles of 563 CpG, while the other employs the methylation profiles of 540 CpGs to estimate *relative* age, which is the ratio of chronological age of an animal to the maximum lifespan of its species; with resulting values between 0 and 1. This relative age ratio is highly advantageous because it allows alignment and biologically meaningful comparison between species with very different lifespans such as cat and human, which cannot otherwise be afforded by direct comparison of their chronological ages.

To arrive at unbiased estimates of the epigenetic clocks, we carried out cross-validation analysis of the training data. To develop the basic cat clock, the training data employed consisted of cat blood DNA methylation profiles, while human and cat DNA methylation profiles constituted the training data for both the human-cat clocks. The cross-validation study reports unbiased estimates of the age correlation R (defined as Pearson correlation between the age estimate – DNAm age – and chronological age), as well as the median absolute error. As indicated by its name, the cat blood clock is highly accurate in age estimation of blood (R=0.97 and median absolute error 0.83 years, **Figure 2A**). The human-cat clock for chronological age is highly accurate when DNA methylation profiles of both species are analyzed together (R=0.98, **Figure 2B**), and remains remarkably accurate when restricted to cat blood samples (R=0.97, **Figure 2C**). Similarly, the human-cat clock for *relative age* exhibits high correlation regardless of whether the analysis is applied to samples from both species (R=0.98, **Figure 2D**) or only to cat samples (R=0.97, **Figure 2E**). This demonstrates that relative age circumvents the skewing that is inherent when chronological age of species with very different lifespans is measured using a single formula.

**Figure 2:**
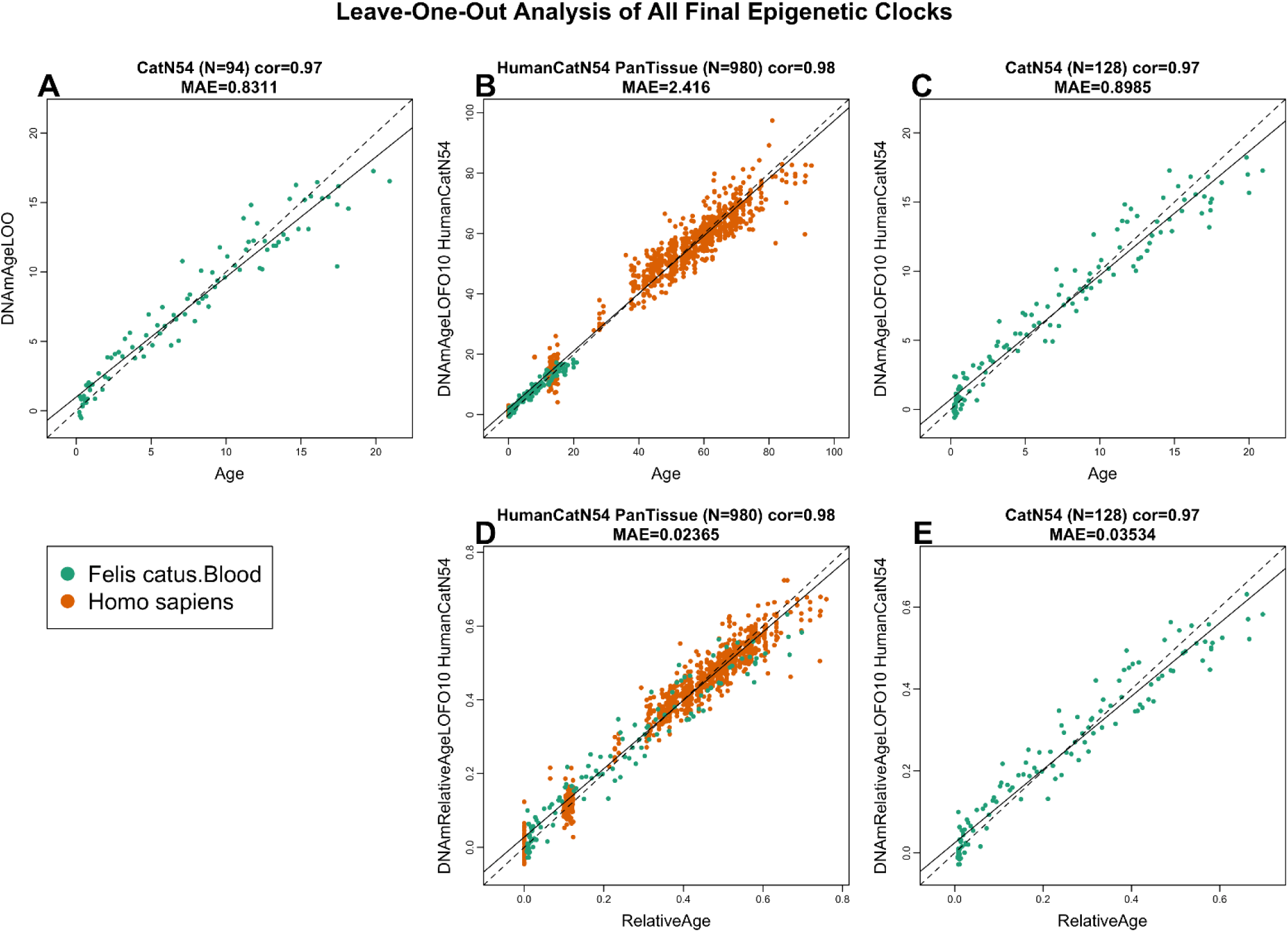
Cross-validation study of epigenetic clocks for domestic cats and humans. A) Epigenetic clock for blood samples from cat. Leave-one-sample-out (LOO) estimate of DNA methylation age (y-axis, in units of years) versus chronological age. B) Ten-fold cross validation (LOFO10) analysis of the human-cat clock for chronological age. Dots are colored by species (green=cat). C) Same as panel B) but restricted to cats. D) Ten-fold cross validation analysis of the human-cat clock for relative age, which is the ratio of chronological age to the maximum recorded lifespan of the respective species. E) Same as panel D) but restricted to cats. Each panel reports sample size, correlation coefficient, median absolute error (MAE).

A cross validation analysis reveals that both human-cat clocks lead to highly accurate estimates in *human* blood and skin samples (R>=0.96, **Figure 3**).

**Figure 3.**
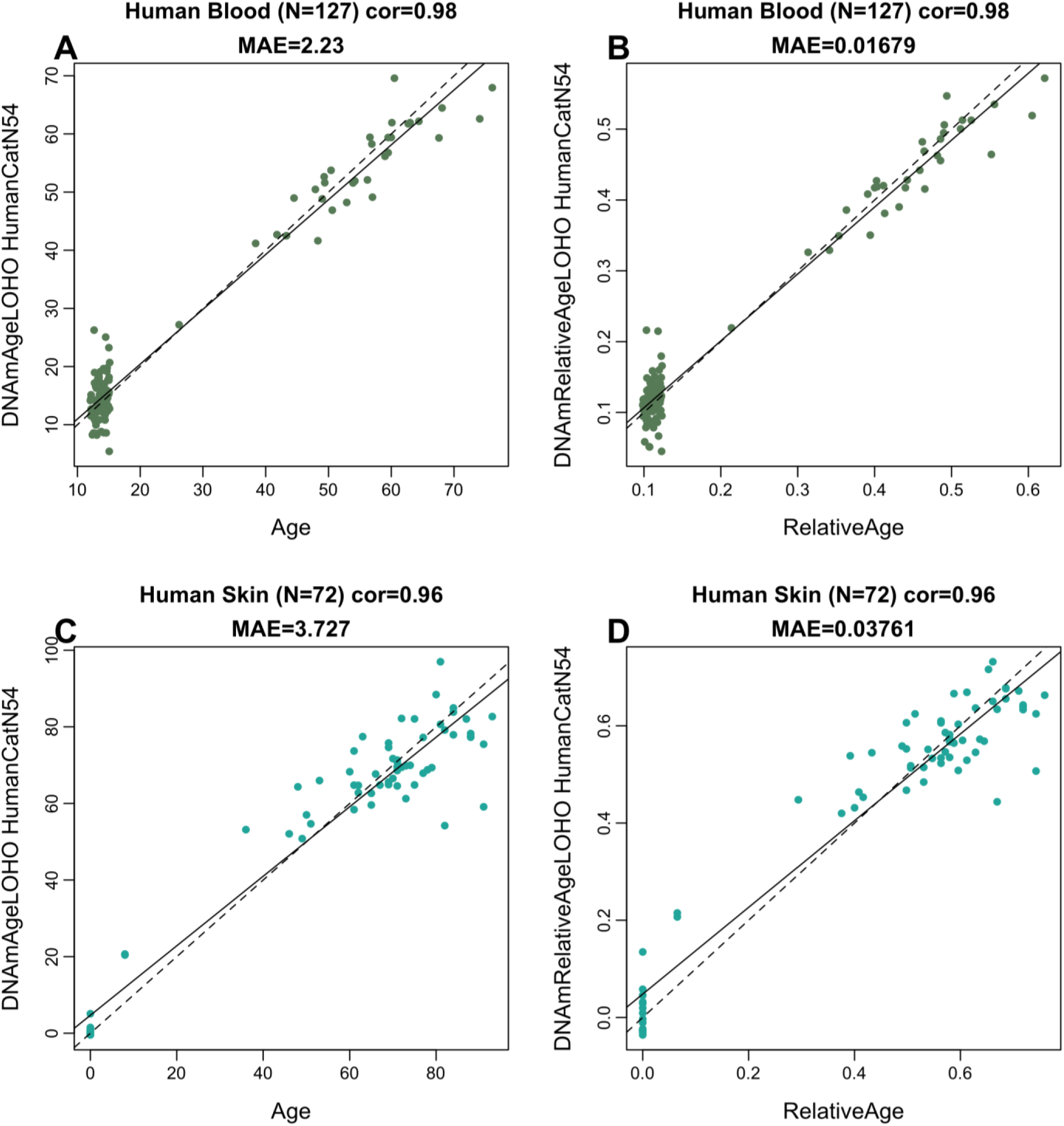
Cross-validation study of human-cat clocks applied to human tissue samples. The y-axis reports leave-one-human sample-out (LOHO) analysis results for the human-cat clocks of chronological age (A,C) and relative age (B,D). Each dot in the upper (A,B) and lower panels (C,D) corresponds to a human blood and skin sample, respectively. Each title reports the sample size, Pearson correlation coefficient, and median absolute error.

## Age-related CpGs

In total, 34851 probes from HorvathMammalMethylChip40 are aligned to loci that are proximal to 5379 genes in *Felis_catus*_9.0.100 genome assembly. Due to the high inter-species conservation of the probes on the array, findings from the cat methylation data can probably be extrapolated to human and other mammalian species. Epigenome-wide association analysis of chronological age revealed a very significant impact of age on DNAm changes (**Figure 4A**). The methylation level of 1379 CpGs altered in function of age, with a significance of p<10^−8^. The CpGs with the greatest methylation changes, and the corresponding proximal genes are as follows: *SLC12A5* promoter (correlation test Z statistic z = 20), *HECTD2* exon (z = -17), hypermethylation in 8 CpGs in *NEUROD1* promoter (z = 8.3 to 16.7), and hypermethylation in two CpGs in *FOXG1* intron (z = 8.9 to 16.4), and 5 CpGs in *FOXG1* exon (z = 8.5 to 11.1, **Figure 4A**). Aging-associated CpGs were distributed in genic, as well as intergenic regions that can be defined relative to transcriptional start sites (**Figure 4B**). In promoters and 5’UTRs, 76% of CpGs increased methylation with age. These regions contain most of the CpG islands, which support our next analysis that showed CpG islands to have a higher positive correlation with age compared to other CpG sites (**Figure 4C**).

**Figure 4.**
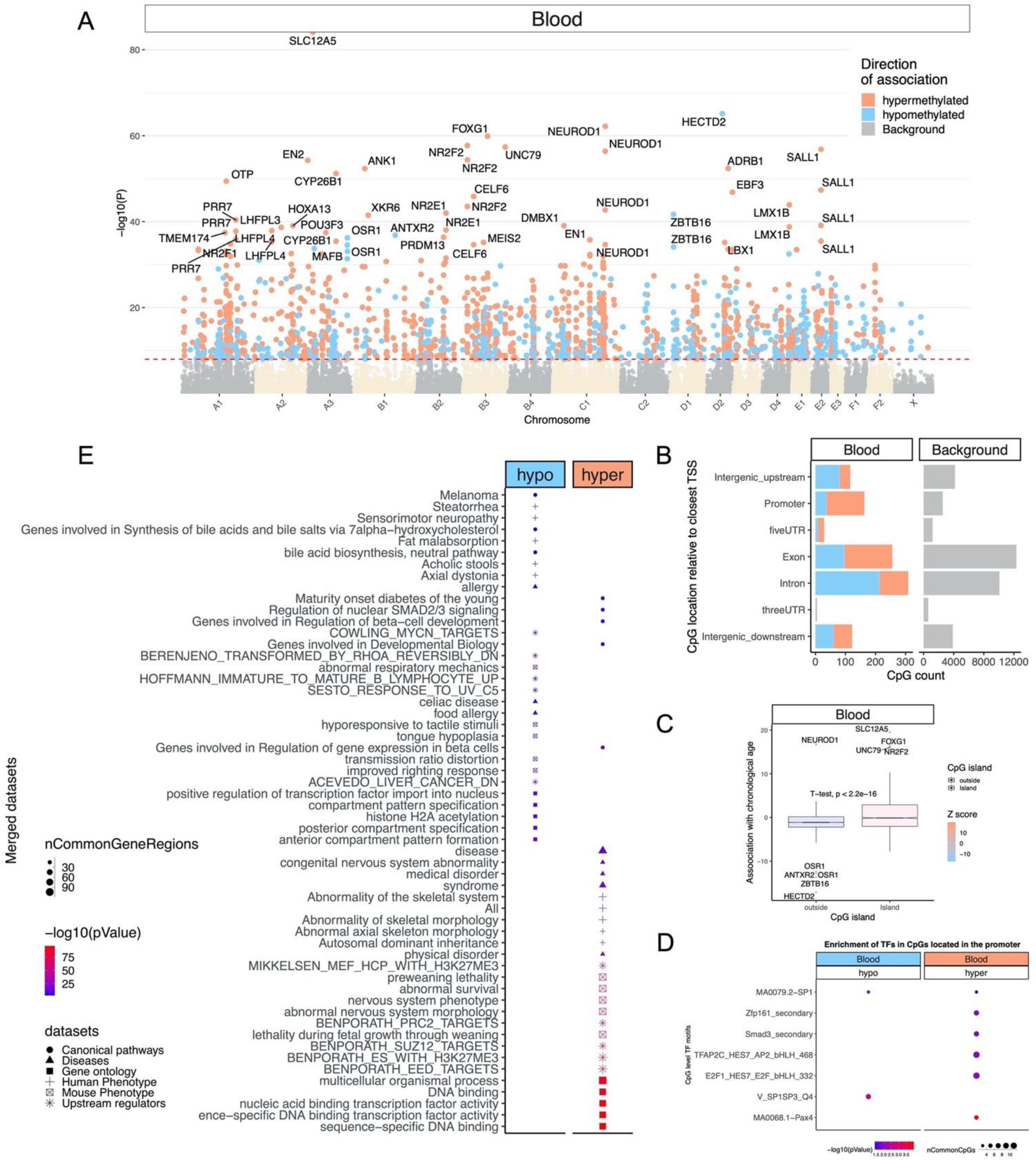
Epigenome-wide association (EWAS) of chronological age in blood of *Felis catus*. A) Manhattan plots of the EWAS of chronological age. The coordinates are estimated based on the alignment of Mammalian array probes to *Felis*_*catus*_9.0.100 genome assembly. The direction of associations with p < 10^−8^ (red dotted line) is highlighted by red (hypermethylated) and blue (hypomethylated) colors. Top 30 CpGs are indicated by their neighboring genes. B) Location of top CpGs in each tissue relative to the closest transcriptional start site. Top CpGs were selected at p < 10^−8^ and further filtering based on z score of association with chronological age for up to 500 in a positive or negative direction. The grey color in the last panel represents the location of 34851 mammalian BeadChip array probes mapped to *Felis*_*catus*_9.0.100 genome. C) CpG islands have higher positive association with age (hypermethylation) than other sites. D) Transcriptional motif enrichment for the top CpGs in the promoter and 5’UTR of the neighboring genes. The motifs were predicted using the MEME motif discovery algorithm, and the enrichment was tested using a hypergeometric test ^50^. E) Enrichment analysis of the top CpGs with positive (hypermethylated) and negative age correlations in feline blood. The gene level enrichment analysis was carried out using the GREAT software ^51^. The background probes were limited to the 25040 probes that could be mapped to the same gene in the cat genome and the human genome (Hg19). The top 3 enriched datasets from each category (Canonical pathways, diseases, gene ontology, human and mouse phenotypes, and upstream regulators) were selected and further filtered for significance at p < 10^−4^.

Transcriptional factor enrichment analysis suggests that hypomethylation in SP1 and hypermethylation in PAX4 binding sites are among the top motifs that show age-associated changes in cat blood (**Figure 4D**). In principle, this would indicate greater access of SP1 protein to some of its binding sites, with increasing age. However, the outcome of this is difficult to predict as SP1 activates the transcription of many genes that are involved in diverse cellular processes ranging from cell growth, apoptosis, immune response to chromatin remodeling. However, the collective enrichment of gene targets of four of the transcription factors together (SMAD3, SP1, SP3, and E2F1) points to their involvement in telomerase regulation (enrichment p =3e-9). In addition, SP1 and E2F1 target genes are also involved in mitophagy (enrichment p =2e-4). The involvement of both, telomerase and mitophagy in ageing, is well-attested in the literature ^36^. PAX4 on the other hand is a transcription factor whose target genes are involved in differentiation and development; likewise, with TFAP2. Gene level enrichment analysis of the significant CpGs highlighted changes in transcription factor activity, development, nervous system changes, and also pathways related to diabetes onset, which all overlap with aging biology in humans and other species (**Figure 4E**). Several potential upstream regulators were also identified as discussed below.

We further examined the enrichment of tissue-type-specific epigenome states for DNAm aging in cats. In both, chromatin state analysis and histone 3 marks, the top tissue type predicted for both hypermethylated and hypomethylated CpGs was blood, which is as it should be, as this is indeed the tissue from which the cat DNA was derived (**Supplementary Figure 2**). The age-associated hypomethylated CpGs were primarily flanking active transcriptional start sites (TSS) and enhancer regions. These CpGs are also marked with H3K4me1 and H3K4me3 modifications which are associated with active transcription. In contrast, age-associated hypermethylation occurs mainly in bivalent/poised TSS, flanking bivalent TSS/Enhancer, bivalent enhancer, and repressed Polycomb binding sites. The histone marks for hypermethylated CpGs included H3K27me27, H3K4me1, and H3K9me3. Collectively, these are all consistent with repression of gene expression from these sites. DNaseI hypersensitive marks (DHS) did not identify blood as our target tissue type, suggesting that age-related DNA methylation changes in nucleosome-depleted (open chromatin) sites is probably not tissue-specific.

### Applying the cat clocks to other species

Having developed these three cat clocks, we used them to estimate the ages of blood DNA methylation profiles from four other mammalian species: guinea pigs, rabbits, ferrets, and alpacas (**Figure 5**). It was not expected that these clocks would estimate their ages accurately. Instead, this was carried out to ascertain the degree by which these cat clocks can predict the age of animals within a non-cat species; *relative to each other*, as indicated by a high correlation coefficient between age and DNAmAge. Understandably, the cat clock and the human-cat clock, which operate with chronological age, registered estimates that are very distant from the chronological age of the animals. Despite this, these two clocks correctly predicted the ages of these animals *relative to each other* (within the same species). This was similarly observed with the second dual species clock, the human-cat relative age clock. It is acknowledged that the paucity of samples, especially of guinea-pig and ferret necessitate some caution in interpretation, but collectively, these results are consistent with the fact that epigenetic clocks developed for one mammalian species can be employed to a limited extent, to other species, and reveal the association of DNA methylation changes with age.

**Figure 5.**
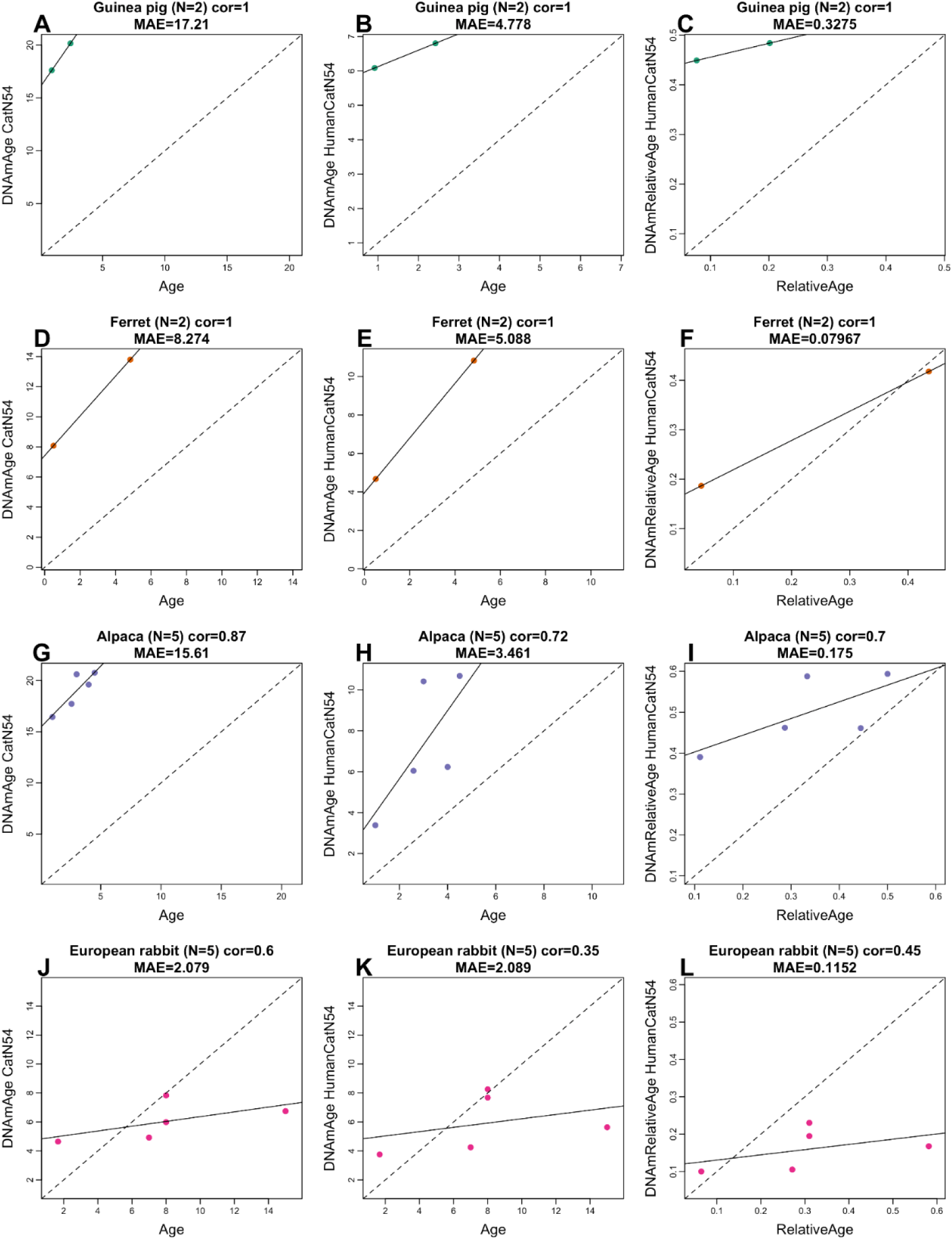
Epigenetic clocks for cats applied to other species. The rows correspond to different species A-C) Guinea pig, D-F) ferret, G-I) alpaca, J-L) European rabbit. Columns correspond to the three epigenetic clocks: A,D,G,J) pure cat clock age estimate (y-axis in years), B,E,H,K) human-cat clock of chronological age in years. C,F,I,L) human-cat clock of relative age. Each panel reports the sample size (N), Pearson correlation, and median absolute error (MAE).

## Discussion

We have previously developed several human epigenetic clocks from DNA methylation profiles that were derived from various versions of human Illumina DNA methylation arrays. As these arrays are specific to the human genome, a critical step toward crossing the species barrier was the bespoke development and use of a mammalian DNA methylation array that profiles 36 thousand CpGs with flanking DNA sequence that are highly conserved across numerous mammalian species. The employment of this array to profile 128 blood samples, represent the most comprehensive epigenetic dataset of domestic cats thus far. These data allowed us to construct highly accurate DNA methylation-based age estimators for the domestic cat that is applicable to their entire life course (from birth to old age). The successful derivation of a cat clock using CpGs that are embedded within evolutionarily conserved DNA sequences across the mammalian class, further confirms the conservation of the biological mechanism that underpins ageing. The ability of the cat clocks to correctly predict ages of animals of the same species (guinea pigs, rabbits, ferrets, and alpacas), *relative to each other* goes a little further in support of this notion. While the mechanism of epigenetic ageing remains to be identified and described, its presence in a host of mammalian species and possibly beyond, indicates an ancient provenance. The potential that the cat clock would contribute to feline health is encouraged by the fact that human epigenetic age acceleration is associated with a wide array of primary traits, health states and pathologies. While it is still unclear why age acceleration is connected to these characteristics, it does nevertheless suggest that extension of similar studies to cats may allow for the development of epigenetic age acceleration as a surrogate or indicator of feline biological fitness; for which there is presently no objective measure.

An equally important potential of the cat clock is the feasibility of including domestic cats in ageing research. The ability to objectively measure biological age is a pre-requisite for such studies and as this is now afforded to cats, the advantage of their inclusion can now be harnessed. Domestic cats share the same living environment as their human owners, but with a lifespan that is considerably shorter. This allows not only investigation into age-affecting factors and potential mitigators of ageing; but also, the impact of these on longevity, which is not readily carried out with humans. However, to accurately translate age-related findings from cats to humans require a correct and accurate measure of age-equivalence. The present rule of thumb where a 1 year-old cat is equivalent to a 15 year-old human, and a 2 years-old cat is equivalent to a 24 years-old human; followed by the addition of 4 years to ever year of a cats life from then on, is a very crude approximation.

We fulfilled this need through a two-step process. First, we combined DNA methylation profiles of cats and humans to generate a dual-species clock (human-cat), which is as accurate in estimating cat age as it is for human age; in chronological unit of years. This demonstrates the feasibility of building epigenetic clocks for different species based on a single mathematical formula. That this single formula is equally applicable to both species effectively demonstrates that epigenetic aging mechanisms are highly conserved. However, the incorporation of two species with very different maximum lifespans, such as cat and human, into a single representative graph, raises the inevitable challenge of polarised distribution of data points along the age range. Furthermore, this does not resolve the challenge of age-equivalence between these two species. We addressed these two challenges simultaneously by expressing the ages of every cat and human in respect to the maximum recorded ages of their respective species (species lifespan), i.e. 30 years for cats and 122 years for humans. The mathematical operation of generating a ratio eliminates chronological unit of time and produces a value that indicates the age of the organism in respect to the maximum age of its own species. This allows a meaningful and realistic cross-species comparison of biological age. For example, the biological fitness of a 20 year-old cat, which is very old, is not equivalent to that of a 20 year-old human, who is young. However, a cat with a relative epigenetic age of 0.5 is more comparable to a human of similar relative epigenetic age. Collectively, the ability to use a single mathematical formula to measure epigenetic age of different species and the replacement of chronological unit of time with proportion of lifespan, are two significant innovations that will propel cross-species research as well as cross-species benefits.

Detailed analyses of age-related methylation changes in cat blood reveals that CpGs that became increasingly methylated with age are located largely within promoters, CpG islands and exons. On the other hand, CpGs that become de-methylated with age are most often found to reside in introns. The consequence of DNA methylation, especially within CpG islands and promoters, is largely suppression of transcription. However, the outcome of intron demethylation is less easy to generalize. Much can be speculated about how these methylation changes can coordinate expression of genes proximal (or perhaps distal too) to these CpGs in function of age. Lack of precise understanding notwithstanding, these epigenetic changes are nevertheless consistent with, and likely mediate, the widely-observed age-related changes in gene expression.

In the absence of empirical data on age-related gene expression changes in cats, we identified genes that are in proximity to age-related CpGs, and then went on to ascertain the intracellular pathways, or diseases/conditions that are associated with these genes. Unsurprisingly, these did not produce results that were immediately obvious and easily understood as being the cause of ageing. Indeed, as our understanding of ageing is at its infancy, separating the cause from the consequence of these age-related changes is a formidable endeavor. Instead, what these results provide is an early glimpse into potential pathways that should be further investigated and tested to ascertain how and why their alterations accompany increasing age. In this regard, it is noteworthy that pathways involved in organism development and maintenance of organ and tissue function constitute a significant portion of the top pathways identified. This contrasts with cancers for example, where alterations to expression of oncogenes, tumour suppressor genes, DNA repair proteins and checkpoints proteins are often encountered. In other words, age-related changes appear to involve development and maintenance of cell function and identity, rather than cellular proliferation or repair. This is consistent with the high score of age-related methylation changes to targets of PAX4 and TFAP2 transcription factors, whose target genes participate in differentiation and development. This is further typified by the fact that the target loci of PRC2, Suz12 and Histone H3K27me3 are identified as being hyper-methylated with age. Suz12 is a component of PRC2, which methylates histone H3K27, which in turn binds to chromatin to prevent transcription of genes that are primarily involved in cell commitment, cellular differentiation and maintenance of cell identity. Interestingly, Suz12 and Histone3K27me3 targets are similarly modified with age, in dogs ^12^. Indeed, such age-related methylation was also previously identified in humans, to occur disproportionately at CpGs in PRC2 target sites ^14^; reinforcing the importance of the process of development in ageing and cross-species conservation of the ageing process. This is further supported by the fact that hypermethylation of bivalent chromatin domain, PRC-binding sites and H3K27me3 featured very strongly with age-related feline CpGs that were analysed with eForge version 2 ^37^. Cross-species concordance of transcription factors, chromatin states, genes and pathways that score highly in computational analyses of age-associated CpGs is a very effective way to identify the most relevant ones from a large number of hits. In this regard, TFAP2, ZFP161 and E2F1/3 are proteins whose binding sites on DNA become increasingly methylated with age in cats as well as bats (manuscript submitted separately). The fact that these far-removed species exhibit these similarities encourages greater attention to be paid to these proteins and their functions. As mentioned above TFAP2 is a transcription factor whose target genes are involved in the cellular differentiation and organ development. ZFP161 protein binds to GC-rich DNA regions, regulates DNA replication fork stability and maintains genomic stability ^38^. It is interesting that another genomic stabilizing protein, the retinoblastoma protein (RB) exerts it effect through binding the E2F transcription factor ^39,40^, whose binding sites become increasingly methylated with age in these two species. This is particularly relevant as genomic instability is a hallmark of cancers ^41^ as well as ageing ^42^. The relationship between these two biological conditions has long been recognised, and their co-appearance in these studies consolidate this relationship and suggest potential common mechanisms that await elucidation. Moreover, the identification of age-associated methylation changes to targets of transcriptional factors that regulate telomerase expression and mitophagy also speaks to the likelihood of this connection. It is notable that regulation of telomerase, mitophagy, genomic instability and epigenetics are implicated in feline ageing, as these are four of the nine identified hallmarks of ageing ^36^. As epigenetic clocks for increasing number of mammals are developed and become available, it will be highly informative to ascertain whether the age-related features (development, and genomic instability) identified in cats and other mammals thus far, would continue to emerge. The emerging picture thus far consolidates the notion that understanding of ageing in mammals such as cats, who intimately share our living environment, can be translated to human ageing. The HorvathMammalMethylChip40, the dual-species (human-cat) clocks, coupled with the concept of relative age that we described here, are innovations that will greatly assist in this endeavor.

## Materials and Methods

### Study samples

#### Feline and other animal blood samples

The DNA archive of the Royal Veterinary College (RVC) was searched for feline ethylenediaminetetraacetic acid (EDTA) blood samples that were residuals from previous routine hematology testing. Cats were selected to represent the widest age range possible based on the available samples with a uniform distribution across the entire range, available breeds and neutering status. As the samples originated from cats that were presented for veterinary investigation, cats were selected to have no or minimal abnormalities on available laboratory data (hematology, serum biochemistry, endocrinology), reviewed by a board certified veterinary clinical pathologists (BSz). The DNA samples were maintained frozen at -80 **°**C for various amount of time (0-11 years). Samples from guinea pigs, rabbits, ferrets, and alpacas were also residual samples from routine patients presented for veterinary care. Sample collection was approved by the Clinical Research Ethical Review Board of the RVC (URN: 2019 1947-2). Genomic DNA from cat blood was extracted using the Zymo DNA extraction kit according to the manufacturer’s instructions. DNA was eluted in water and quantified with picogreen kit according to the instructions provided.

#### Human tissue samples

To build the human-cat clock, we analyzed previously generated methylation data from n=850 human tissue samples (adipose, blood, bone marrow, dermis, epidermis, heart, keratinocytes, fibroblasts, kidney, liver, lung, lymph node, muscle, pituitary, skin, spleen) from individuals whose ages ranged from 0 to 93. The tissue samples came from three sources. Tissue and organ samples from the National NeuroAIDS Tissue Consortium ^43^. Blood samples from the Cape Town Adolescent Antiretroviral Cohort study ^44^. Skin and other primary cells provided by Kenneth Raj ^45^. Ethics approval (IRB#15-001454, IRB#16-000471, IRB#18-000315, IRB#16-002028).

### DNA methylation data

We generated DNA methylation data using the custom Illumina chip “HorvathMammalMethylChip40”. The mammalian methylation array is attractive because it provides very high coverage (over thousand X) of highly conserved CpGs in mammals but it focuses on only 36k CpGs that are highly-conserved across mammals. DNA methylation arrays were profiled using a custom Illumina methylation array (HorvathMammalMethylChip40) based on 38k CpG sites in highly conserved regions in mammals.

Two thousand out of 38k probes were selected based on their utility for human biomarker studies: these CpGs, which were previously implemented in human Illumina Infinium arrays (EPIC, 450K) were selected due to their relevance for estimating age, blood cell counts, or the proportion of neurons in brain tissue. The remaining 35,988 probes were chosen to assess cytosine DNA methylation levels in mammalian species. Toward this end, an algorithm was employed to identify highly conserved CpGs across 50 mammalian species: 33,493 Infinium II probes and 2,496 Infinium I probes. Not all probes on the array are expected to work for all species, but rather each probe is designed to cover a certain subset of species, such that overall all species have a high number of probes. The particular subset of species for each probe is provided in the chip manifest file. The SeSaMe normalization method was used to define beta values for each probe ^46^.

### Penalized Regression models

Details on the clocks (CpGs, genome coordinates) and R software code are provided in the Supplement. Penalized regression models were created with glmnet ^47^. We investigated models produced by both “elastic net”regression (alpha=0.5). The optimal penalty parameters in all cases were determined automatically by using a 10 fold internal cross-validation (cv.glmnet) on the training set. By definition, the alpha value for the elastic net regression was set to 0.5 (midpoint between Ridge and Lasso type regression) and was not optimized for model performance.

We performed a cross-validation scheme for arriving at unbiased (or at least less biased) estimates of the accuracy of the different DNAm based age estimators. One type consisted of leaving out a single sample (LOOCV) from the regression, predicting an age for that sample, and iterating over all samples. A critical step is the transformation of chronological age (the dependent variable). While no transformation was used for the blood clock for cats, we did use a log linear transformation for the dual species clock of chronological age (**Supplement**).

### Relative age estimation

To introduce biological meaning into age estimates of cats and humans that have very different lifespan; as well as to overcome the inevitable skewing due to unequal distribution of data points from cats and humans across age range, relative age estimation was made using the formula: Relative age= Age/maxLifespan where the maximum lifespan for the two species was chosen from the anAge data base ^48^.

### Epigenome wide association studies of age

EWAS was performed in each tissue separately using the R function “standardScreeningNumericTrait”from the “WGCNA”R package^49^. Next the results were combined across tissues using Stouffer’s meta-analysis method.

## URLs

## Acknowledgements

This work was supported by the Paul G. Allen Frontiers Group (SH). National Institute of Aging 1U19AG057758

## SUPPLEMENTARY MATERIAL

**Supplementary Figure 1.**
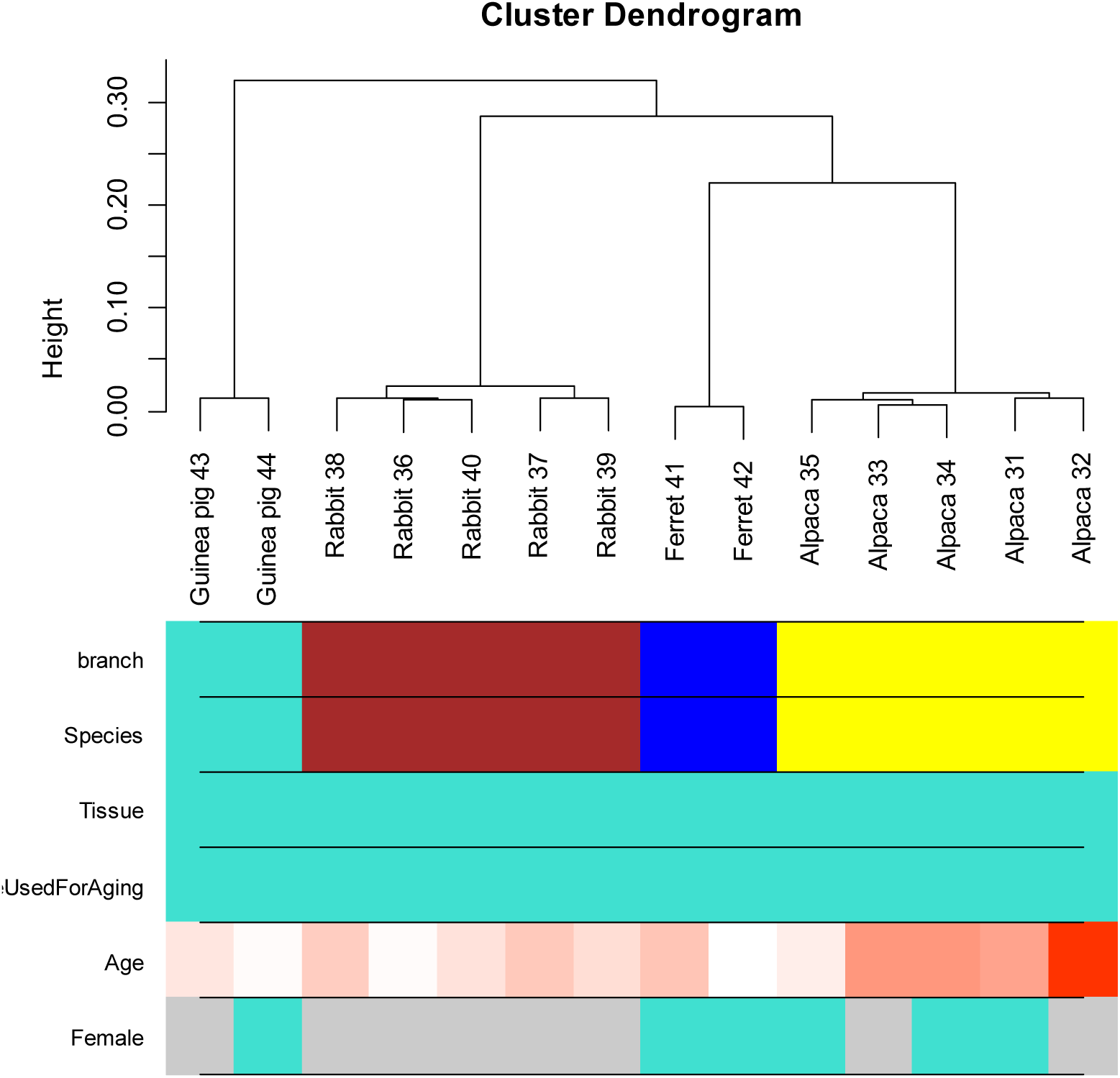
Unsupervised hierarchical clustering of blood samples from guinea pigs, rabbits, ferrets, and alpacas. Average linkage hierarchical clustering based on the inter-array correlation coefficient (Pearson correlation). The first color band, cluster branch, corresponds to a height cut-off of 0.04. The second color band shows that the blood samples cluster by species.

**Supplementary Figure 2.**
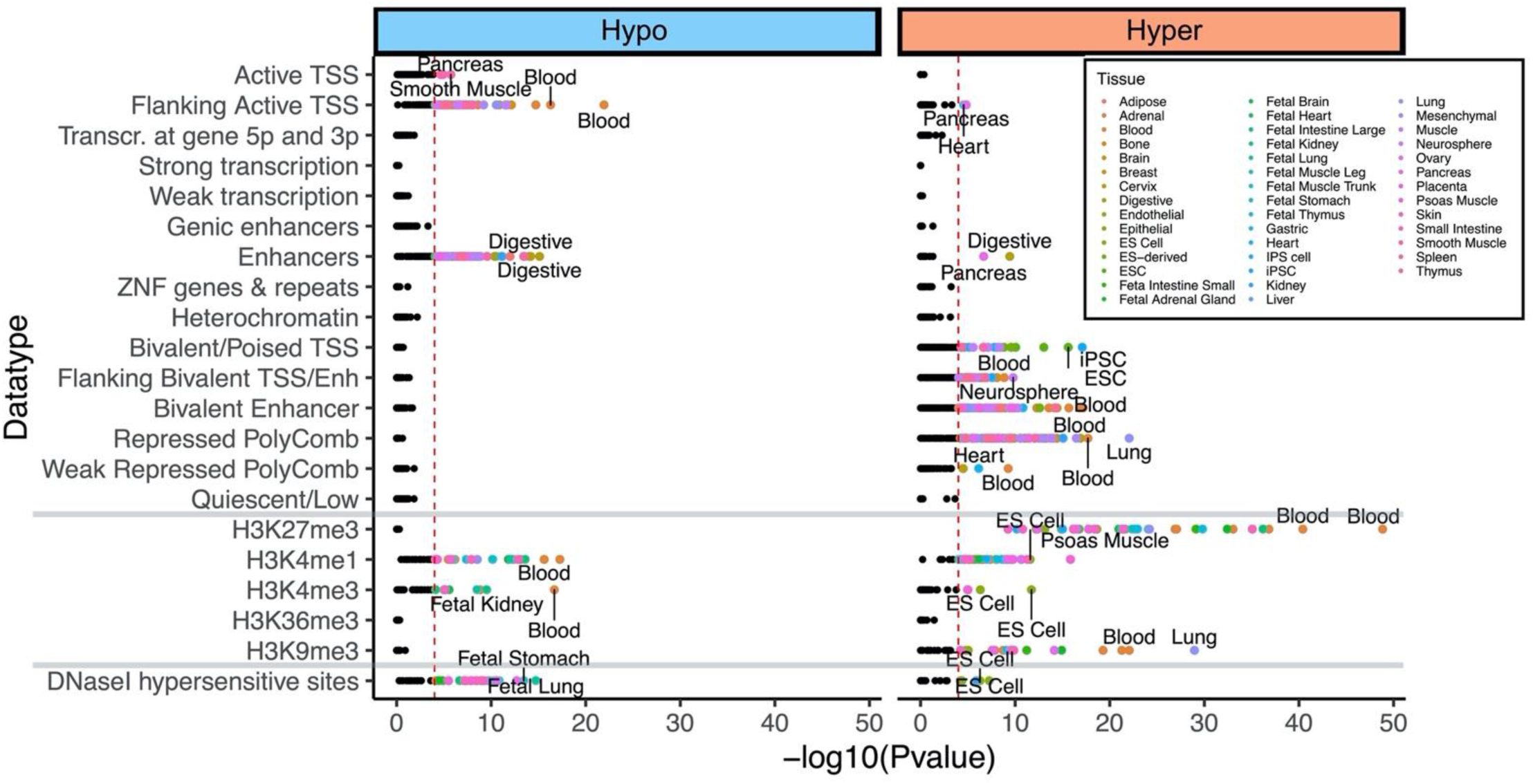
Chromatin state enrichment analysis of age related CpGs with eForge. We analyzed 15 chromatin states, histone 3 marks, and DNaseI I hypersensitivity sites for age-associated CpGs in cats. Highlighted points indicates p < 10^−4^. Top two tissue types for each significant mark are labeled. We selected the *Felis catus* genome as background in eForge V2.0 ^37^.

